# Advancements in Fire-Related Toxic Gas Detection and Prophylactic Strategies: A Focus on Cyanide Concentration Analysis and Antidote Efficacy in Controlled Smoke Inhalation Models

**DOI:** 10.1101/2025.09.21.677646

**Authors:** Jowy Tani, Jia-Long Chen, Wei-Chuan Liao, Chia-Hung Chen, Chau-Hui Wang, Tzu-hui Sun, Yow-Ling Shiue, Jin-Wu Tsai

## Abstract

Between 2020 and 2023, rising global temperatures and extreme weather events have significantly increased fire incidents, releasing toxic smoke containing lethal cyanide gas. Cyanide disrupts cellular respiration and poses severe risks to victims and first responders. This study quantifies cyanide concentrations in fire smoke and evaluates the antidotal efficacy of hydroxocobalamin. Using a smoke chamber calibrated to 100 ppm cyanide, treated mice demonstrated a 70% survival rate compared to 30% in controls, with median survival times of 12 and 5 minutes, respectively (p < 0.05). Employing UPLC-MS and colorimetric analysis, cyanide levels were precisely measured. Furthermore, aerosolized hydroxocobalamin proved effective for rapid absorption. These findings highlight advanced antidotal strategies as pivotal in enhancing survival and shaping fire smoke safety protocols.

## Introduction

In addition to rising global temperatures and increasing extreme weather events, the use of modern electrical equipment has become a significant factor in fire incidents. For example, the proliferation of electric vehicle charging facilities and overloaded circuits have substantially increased the risk of electrical fires. The International Association of Fire and Rescue Services (CTIF) reports that electrical fires account for a notable proportion of global fire incidents, with firefighter fatalities associated with fire events also on the rise [1].

In addition to wildfires, industrial fires are another major source of toxic fire smoke. The combustion of various chemicals used in industrial facilities releases highly toxic gases, posing severe health risks. Cyanide, in particular, is a dangerous component of fire smoke, produced during the combustion of nitrogen-containing materials such as melamine, wool, and silk. Cyanide rapidly inhibits cellular respiration, leading to hypoxia and, if untreated, can be fatal [2].

Smoke inhalation is a significant cause of morbidity and mortality in fire incidents. The gases, particulates, and toxic substances, such as cyanide, present in smoke pose severe threats to fire victims and firefighters. Cyanide is a potent cellular toxin whose rapid inhibition of cellular respiration makes it a critical component of fire smoke toxicity. Particularly in industrial fires, the combustion of melamine can produce extremely high and lethal concentrations of cyanide. At high temperatures, melamine combustion generates cyanide in quantities sufficient to be immediately fatal, posing extreme risks to firefighters entering fire environments [3].

Given the multifaceted issue of fire smoke toxicity, our research aims to explore cyanide concentrations and their acute toxic effects in smoke. By conducting controlled experiments in a smoke chamber with cyanide concentrations set at 100 ppm, we seek to quantify the lethal concentration of cyanide in fire smoke and understand its dynamics in a controlled environment [2, 3].

Cyanide is frequently generated through the combustion of various synthetic materials, including foams like melamine foam. Due to cyanide’s rapid inhibition of cellular respiration, it can cause severe toxic reactions. This study removes the erroneous analogy between “cyanide exposure and vitamin B12 deficiency” and instead focuses on cyanide toxicity and the established mechanisms of vitamin B12, particularly hydroxocobalamin, in detoxification [4].

Beyond assessing cyanide toxicity in smoke, our study also aims to evaluate the efficacy of rescue drugs, such as hydroxocobalamin (Cyanokit), in mitigating cyanide’s toxic effects [5]. Existing literature indicates that hydroxocobalamin is an effective antidote for cyanide poisoning in smoke inhalation scenarios. Our research will contribute to this body of knowledge by comparing the survival rates of treated and untreated subjects in controlled experiments [6,7,8].

Our experiments will use mice as models due to their physiological similarities to humans in toxicology studies. We will record survival rates and times in the smoke chamber to provide valuable data on cyanide’s lethality and antidote effectiveness [3, 4].

Considering the complexity and potential variability of fire smoke composition, our study will employ Ultra Performance Liquid Chromatography-Mass Spectrometry (UPLC-MS) to precisely analyze cyanide concentrations in both water solutions and smoke samples. UPLC-MS is a powerful analytical technique that enables accurate detection and quantification of cyanide, providing a detailed understanding of its presence and concentration in different matrices [9, 10].

The ultimate goal of our research is to provide insights that can inform the development of more effective fire smoke prevention strategies and treatment protocols. By understanding the specific risks associated with cyanide in fire smoke and evaluating the efficacy of existing antidotes, we aim to contribute to a body of knowledge that helps protect civilians and firefighters from the dangers of smoke inhalation.

Additionally, our research plans to explore the potential of aerosolized antidote delivery in fire rescue operations. Aerosolized delivery can rapidly administer the drug to the lungs, enhancing absorption efficiency and providing immediate detoxification effects at the fire scene. This delivery method offers significant advantages for rescue personnel operating in high-risk environments [11,12,13].

These efforts will provide scientific foundations and practical guidance for improving fire smoke prevention and treatment, ultimately enhancing public safety and rescue efficiency.

## Materials and Methods

### Animal Model and Ethical Approval

In our study, we adhere to the highest standards of ethical treatment for the animal subjects used. Male B6(C57BL/6) mice, aged between 8–10 weeks, acquired from LASCO, are chosen for their physiological relevance to the objectives of the study. Each mouse is given a minimum of one week to acclimatize to the laboratory environment prior to the commencement of experiments, ensuring minimal stress and adaptation issues [14].

Humane endpoints were established in accordance with PLOS ONE and IACUC guidelines. Mice were continuously monitored during exposure and throughout the observation period. Animals exhibiting signs of severe distress, including rapid or labored breathing, marked reduction in activity, loss of righting reflex, or unconsciousness, were immediately removed from the experiment and euthanized.

Euthanasia was performed under general anesthesia using isoflurane inhalation, ensuring minimal pain or distress. Isoflurane anesthesia was also administered prior to terminal blood collection and other invasive procedures. Analgesia and refinement measures were implemented to minimize suffering, including close monitoring, acclimatization prior to experiments, and immediate medical intervention if unexpected adverse events occurred.

Post-experiment, all animals are continuously monitored for an additional week to observe any adverse reactions or long-term effects stemming from the experimental procedures. This extended observation period is crucial for identifying any delayed responses to the treatment or exposure, ensuring the welfare and health of the animals throughout the study duration.

Our experimental protocols strictly adhere to the principles of the 3Rs (Replacement, Reduction, and Refinement) in animal research. Approval was obtained from the Institutional Animal Care and Use Committee (IACUC) of National Yang Ming Chiao Tung University, with certification number IACUC1121001.

### Drug Nebulization and Safety Assessment

The combination of these two drugs offers a comprehensive approach to preventing the harmful effects of fire smoke, particularly focusing on cyanide poisoning and free radical damage. The primary target of the nebulized drug combination is the respiratory tract, including the mouth, nose, bronchi, lungs, and deep alveoli. The drugs, when inhaled, provide protection along the entire respiratory pathway. They adhere to the surface of the tissues, ensuring that when toxic gases are inhaled, most of the cyanide is neutralized before entering the bloodstream. This significantly reduces the cyanide’s potential to damage blood cells and impair their oxygen-carrying function [12].

Although the concentration of inhaled drugs in the bloodstream is often low and difficult to detect, we use a dual approach to analyze drug concentrations by collecting both blood and respiratory tissue fluid samples. We employ UPLC-MS which can detect smaller units, to measure drug concentrations accurately [10].

The nebulizer used was HEALTH & LIFE CO., LTD.’s HL-100 model (FDA approved nebulizer, K081738). The drug combination included Hydroxocobalamin (from Tokyo Chemical Industry Co., Ltd.) and Deferoxamine (from Novartis AG), dissolved in a 0.3M Tris-buffer solution (Tris(hydroxymethyl)aminomethane). Post-inhalation, animals were monitored for one week to ensure no adverse allergic reactions or abnormalities were observed (Supple figure1).

### Fire Smoke Experiment and Cyanide Concentration Testing

To simulate a fire smoke environment with high hydrogen cyanide (HCN) concentrations, 0.02–0.03 g of melamine-based foam was combusted in a sealed 500 mL glass bottle. Upon completion of combustion, the bottle was immediately sealed to retain the generated smoke. A conduit was then employed to directly connect the combustion bottle’s opening to the BW Ultra (Honeywell BW) HCN detector for real-time measurement of HCN concentration in ppm. Additionally, to precisely quantify HCN concentration, an aqueous sampling method was simultaneously employed. In this method, the combustion bottle was pre-filled with purified water, and after combustion, the bottle was vigorously shaken to rapidly dissolve HCN gas into the water. The obtained water samples were analyzed using an HCN detection reagent (LU HENG) via colorimetric assay, which relies on the distinct color gradient produced by varying HCN concentrations (Supple figure 5). Concentrations were then determined quantitatively by comparison to a previously established calibration curve for HCN. Based on a fixed gas flow rate and sampling time, the distribution range of the HCN concentration was thus accurately established, ensuring each experiment was conducted within the desired concentration range. Given the high toxicity of HCN, strict safety protocols were observed throughout the experiment by using appropriate personal protective equipment (e.g., gas masks, gloves), conducting all smoke tests within a fume hood, and notifying the environmental safety center in advance. After multiple trials, a stable HCN environment was established, providing a reliable basis for the fire smoke exposure model [15].

Mice were divided into two groups for the fire smoke efficacy test. One group inhaled experimental water as a control, while the other inhaled the drug combination as a preventative measure. In a small-scale fire model, one mouse from each group (control and drug) was exposed to cyanide concentrations controlled between 100-110 ppm (Fig.1, Supple figure 2). The maximum observation time per experiment was 15 minutes, with immediate cessation upon any signs of apparent death or cessation of respiration and heartbeat. The survival time of each mouse was recorded. After completing all group experiments, survival rates and median survival times were statistically analyzed. Surviving mice were monitored for at least one week post-experiment, with immediate medical intervention provided in case of distress.

**Figure 1.**
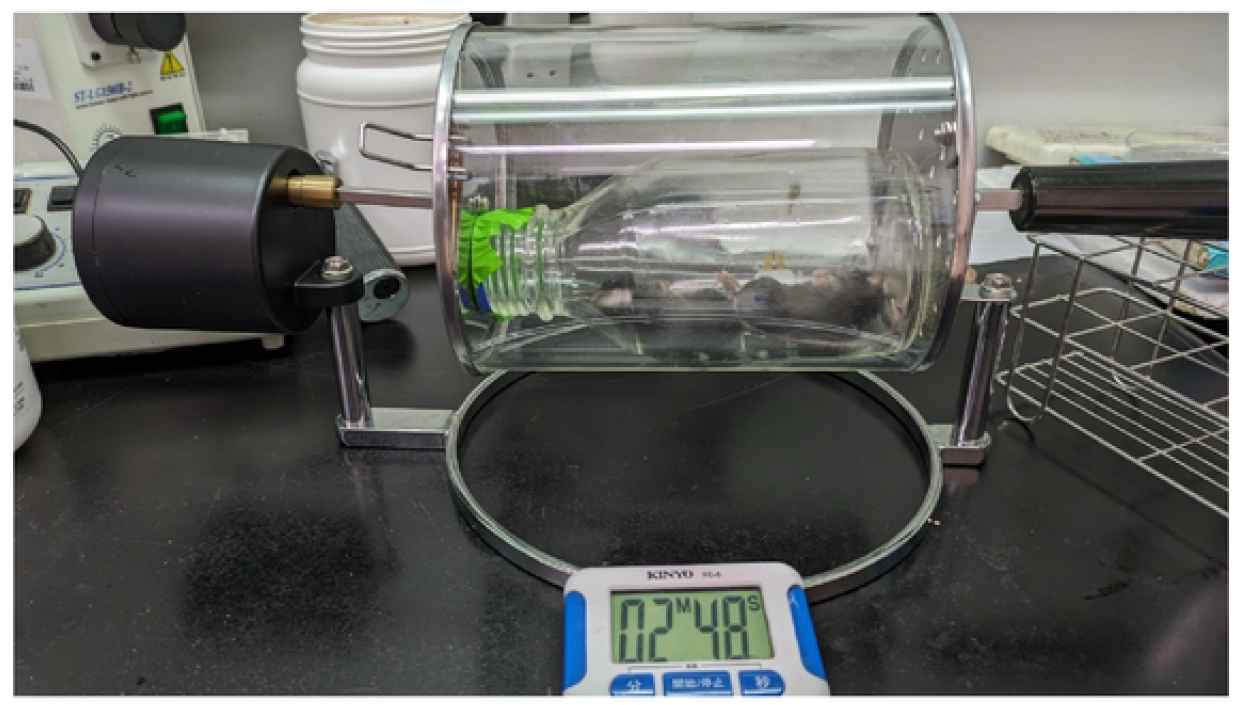
Fire Smoke Model. Mouse exposure setup. A representative image shows a mouse positioned within a sealed exposure chamber where controlled HCN-rich smoke is introduced via the combustion of a small amount of polyurethane foam.

### Drug Formulation

Two formulations of the combination drug were prepared:

1. **Low dose:** Hydroxocobalamin 25 mg + Deferoxamine 1 mg in one dose of 0.3M Tris buffer.
2. **High dose:** Hydroxocobalamin 150 mg + Deferoxamine 6 mg in one dose of 0.3M Tris buffer.

### Nebulization Procedure

Nebulization time for low-dose and high-dose combination drug administration in B6 mice. Mice were exposed to aerosolized Hydroxocobalamin and Deferoxamine under controlled conditions using a nebulizer. All mice were contained within a designated exposure unit to maintain consistent drug delivery conditions [6, 12, 16].

For the low-dose formulation (Hydroxocobalamin 25 mg + Deferoxamine 1 mg in 0.3M Tris buffer), the nebulization time averaged around 1 minute 35 seconds. For the high-dose formulation (Hydroxocobalamin 150 mg + Deferoxamine 6 mg in 0.3M Tris buffer), the nebulization time averaged around 4 minutes 12 seconds. (Supple figure1).

### Sample Collection

Blood and lung fluid samples were collected at specific time points post-administration (20 minutes, 1 hour, and 2 hours). Samples were analyzed to determine concentrations of hydroxocobalamin and cyanocobalamin using UPLC-MS (Table 3) [9,10].

### Sample Analysis Method

In this study, we analyzed both blood and lung fluid samples using UPLC-MS (Ultra Performance Liquid Chromatography-Mass Spectrometry) for metabolomic profiling. The samples were collected, frozen, and stored at −20°C. For preparation, 60 µl of each sample was mixed with 240 µl of methanol to precipitate proteins. The mixture was vortexed, followed by centrifugation at 15,000g for 15 minutes at 4°C. Subsequently, 252 µl of the supernatant was transferred to an Eppendorf tube, vacuum-dried, and the residue reconstituted in 48 µl of water. The reconstituted solution was further centrifuged at 15,000g for 10 minutes at 4°C to remove precipitates. For the final analysis, 10 µl of the processed sample was injected into a Waters Xevo TQD mass spectrometer connected to a Waters ACQUITY UPLC system. Chromatographic separation was carried out on a BEH C18 column, and data was acquired in SRM (Selected Reaction Monitoring) mode with positive electrospray ionization (ESI+) [9,10].

### Enhanced Statistical Analysis Description for Academic Submission

#### Mann-Whitney U Test

Comparison: This test compared the survival times of two groups of mice, each consisting of 12 animals, under conditions of fire smoke exposure. The control group and the drug-treated group were assessed to determine if there was a significant difference in their overall survival times.

Statistical Difference: The Mann-Whitney U test yielded a p-value of 0.04, indicating a statistically significant difference in survival times between the control and drug groups, marked as significant (*p < 0.05).

#### Log-Rank (Mantel-Cox) Test

Comparison: This test was utilized to compare the survival distributions over the 15-minute exposure period between the two groups. It specifically evaluated whether the probability of survival differed significantly over time between the control and drug-treated mice.

Statistical Difference: The results from the Log-Rank test would indicate whether the survival experiences of the two groups are statistically different, shedding light on the drug treatment’s efficacy.

## Results

## Conclusion

In addition to these findings, our study highlights the importance of addressing cyanide exposure risks in fire scenarios. Tailored training and protective equipment for firefighters are essential for managing environments prone to high cyanide concentrations. Future clinical trials should evaluate the feasibility of deploying nebulized antidotes in real-world fire scenarios, focusing on their potential to enhance firefighter safety and emergency medical interventions [6, 7].

Our research confirms the acute toxicity of cyanide in fire smoke and demonstrates the efficacy of hydroxocobalamin and deferoxamine in mitigating these dangers [2, 3].

Through controlled experiments in a smoke chamber, we observed significant differences in survival rates and times between the control group and the drug-treated group. Mice exposed to fire smoke without any protective intervention showed a 30% survival rate and a median survival time of 5 minutes, whereas those treated with hydroxocobalamin and deferoxamine achieved a 70% survival rate and a median survival time of 12 minutes (p < 0.05)(Figure 2, 3, Table 1, Supple figure 2).

**Figure 2.**
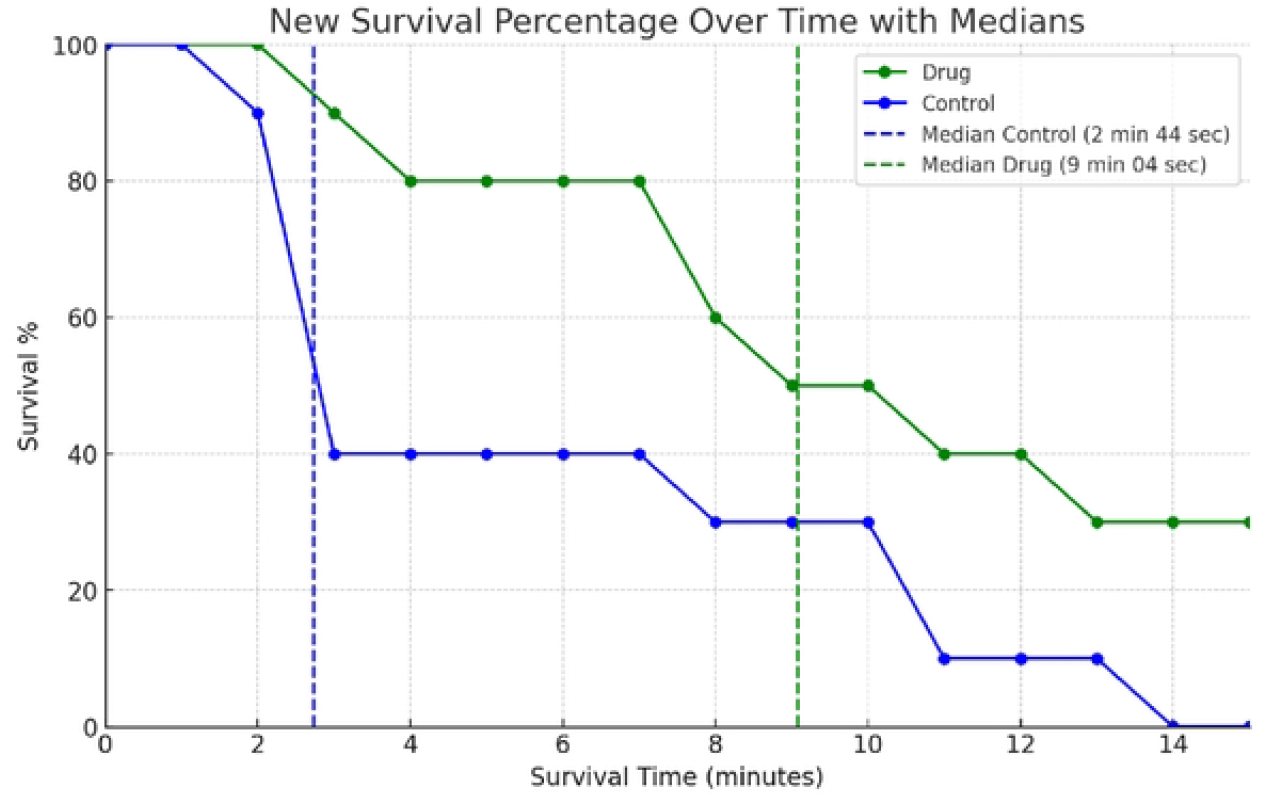
Survival Rate Plot Legend. Survival Rate of Mice Exposed to Fire Smoke for 15 Minutes. This graph illustrates the survival percentage over time for two groups of mice, each consisting of nine individuals, exposed to fire smoke. The control group (blue line) and drug-treated group (green line) were monitored for respiratory cessation or signs of evident mortality, at which point survival time was recorded. The x-axis represents survival time in minutes, capped at 15 minutes, and the y-axis depicts the survival percentage. The vertical dashed lines indicate the median survival times for each group. Control group number=10; Drug prevention number= 10. Drug dose for each group: Hydroxocobalamin 25 mg + Deferoxamine 1 mg in one dose of 0.3M Tris buffer.

**Figure 3.**
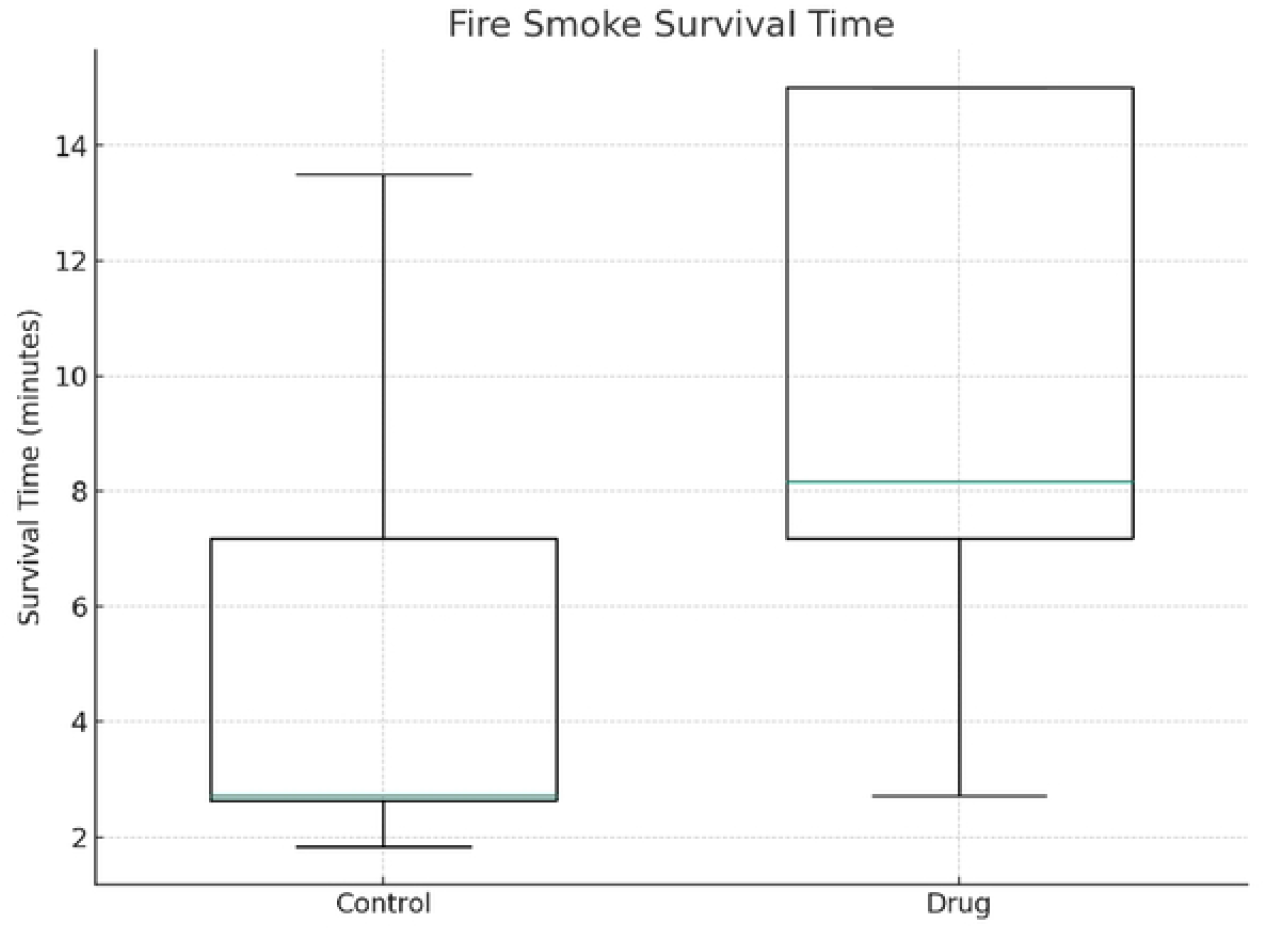
Boxplot Legend. Boxplot Comparing Survival Times of Control and Drug-Treated Mice. This boxplot displays the distribution of survival times for two groups of nine mice each, following a 15-minute exposure to fire smoke. Survival times were recorded at the moment of respiratory cessation or apparent death. The control group is represented in blue, and the drug-treated group in green. The boxplot shows the median, quartiles, and potential outliers in survival times for each group. Mann-Whitney U Test, P value= 0.04

**Table 1.** Mouse Survival Analysis under Varying HCN Concentrations and Exposure Durations.

| **HCN (ppm)** | **Survival Time (Sec)** | **Mice number** | **Dead(Yes/NO)** |
| --- | --- | --- | --- |
| **44** | 900 | 6 | No |
| **74** | 1800 | 2 | No |
| **80** | 1800 | 1 | Yes |
| **80** | 1800 | 1 | No |
| **83** | 600 | 8 | No |
| **101** | 600 | 1 | No |
| **103** | 600 | 1 | No |
| **108** | 780 | 1 | Yes |
| **110** | 167 | 3 | Yes |
| **111** | 149 | 1 | Yes |
| **120** | 182 | 1 | Yes |
| **133** | 137 | 1 | Yes |
| **202** | 110 | 1 | Yes |
| **213** | 130 | 5 | Yes |
| **220** | 71 | 3 | Yes |

Survival outcomes of mice were assessed under controlled exposure to different concentrations of hydrogen cyanide (HCN) and varied exposure durations [3].
. Mice were placed in a controlled environment with specific HCN concentrations, and survival was monitored at intervals to determine the lethal dose and time-dependent effects of HCN exposure. The data are summarized in Table 1, where each row represents a distinct concentration-duration pairing, detailing the percentage of mice surviving at each condition. This analysis provides insights into the acute toxicity profile of HCN, establishing the correlation between concentration, exposure time, and survival probability.

**Table 2.**
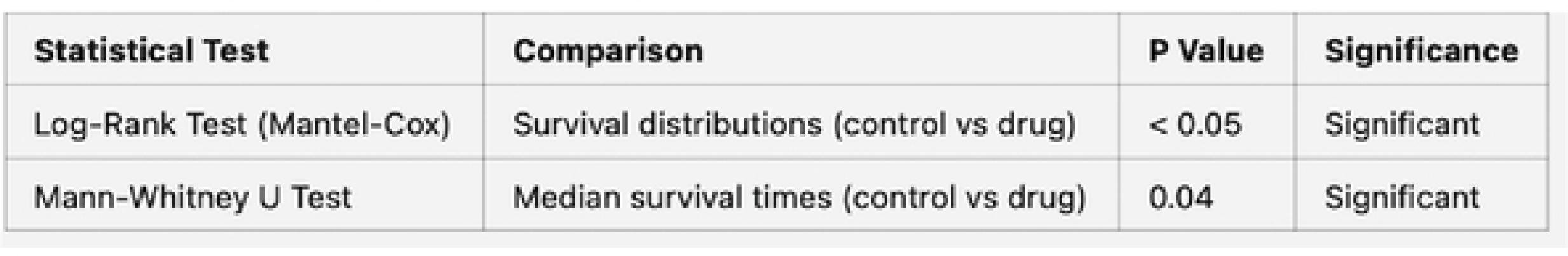
Statistical analyses table.

The analysis of cyanide concentrations revealed consistent levels of 100-110 ppm in the smoke, providing a reliable baseline for assessing toxic exposure. Using advanced analytical methods, including UPLC-MS and colorimetric analysis, we quantified cyanide levels and demonstrated the effectiveness of aerosolized hydroxocobalamin and deferoxamine in reducing toxicity. The aerosolized delivery method proved efficient, with significantly higher drug concentrations detected in lung tissue compared to blood, indicating effective localized delivery (Table 4).

**Table 3.**
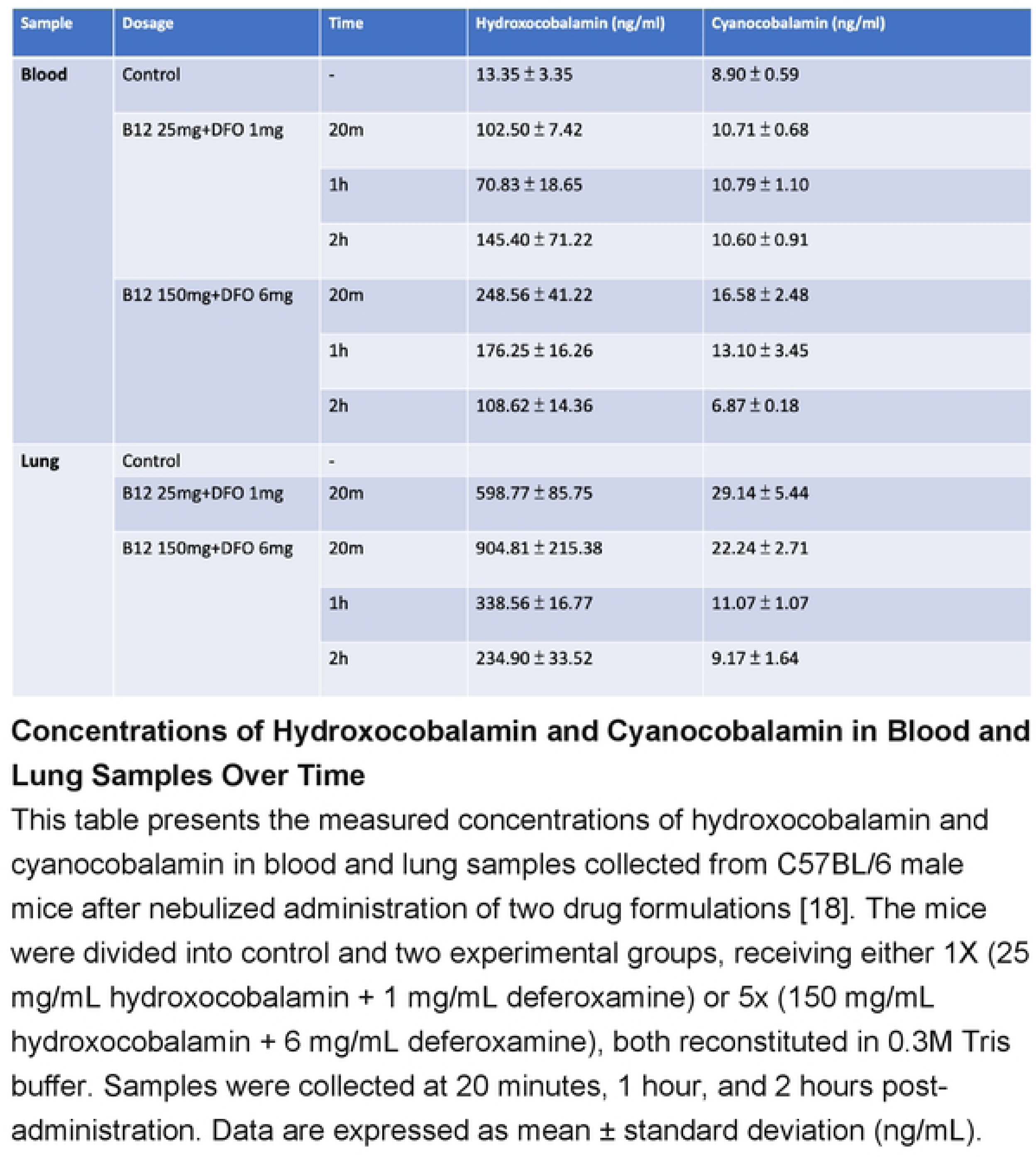
Inhalation Dynamics of Differential Dosages and Concentrations in Fire Smoke Models.

**Table 4.**
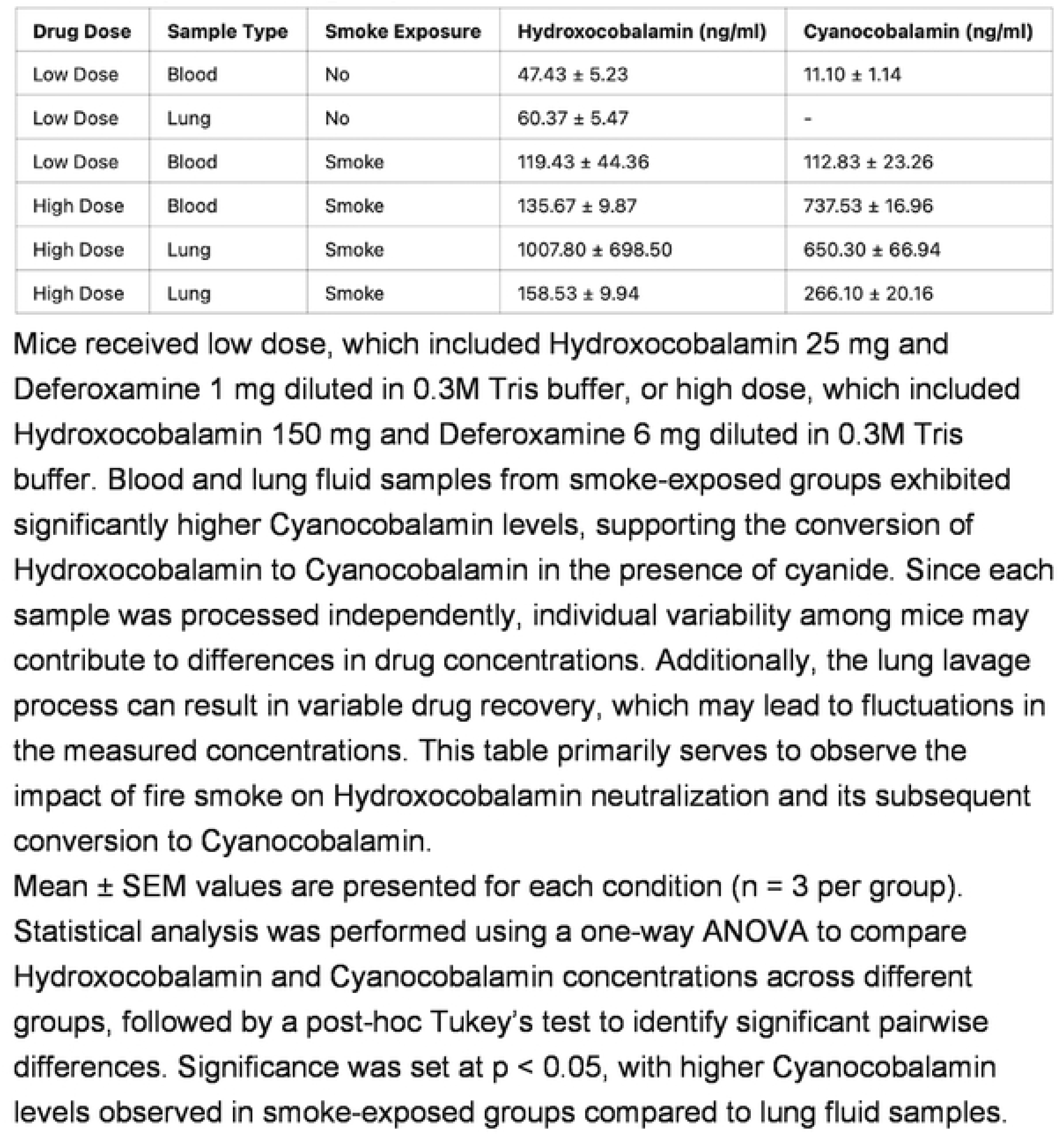
Comparison of Hydroxocobalamin and Cyanocobalamin concentrations in lung fluid (L) and smoke inhalation samples (S) across various experimental groups.

The combination of hydroxocobalamin and deferoxamine offers a dual mechanism: hydroxocobalamin neutralizes cyanide by converting it into cyanocobalamin, while deferoxamine provides protection against oxidative damage caused by fire smoke (Table 3). This comprehensive approach underscores the potential of nebulized antidotes as a practical and effective solution for managing cyanide poisoning during fire incidents. Exposure to smoke induced significant inflammatory responses and tissue damage in lung tissue. Preventive drug administration demonstrated protective effects against smoke-induced lung injury, with dose-dependent improvements observed at higher drug doses (Supple figure 6). Inhalation of the tested drug, even at a high dosage, did not induce any detectable adverse hepatic changes. The histological findings suggest good hepatic safety and tolerability of the drug formulation (Supple figure 7).

While our findings provide strong evidence for the efficacy of these antidotes, further research is needed to address limitations and explore long-term outcomes. This study lays a solid foundation for future clinical research on aerosolized hydroxocobalamin and deferoxamine, aiming to improve survival and safety in real fire scenarios. The insights gained here contribute to advancing fire smoke prevention strategies, shaping public safety policies, and enhancing emergency medical interventions to protect both civilians and firefighters.

## Discussion

Firefighters must be exceptionally prepared for the risk of cyanide poisoning in specific types of fires, including: (1) Residential Fires, (2) High-Rise Building Fires, (3) Industrial and Chemical Plant Fires, (4) Vehicle Fires, (5) Warehouse Fires, (6) Fires in Public Spaces with Synthetic Materials, (7) Transportation and Transit System Fires, and (8) Fires at Plastic Recycling or Disposal Facilities [2, 13].

The results of our study highlight several key insights and implications for the management of cyanide toxicity in fire smoke and the efficacy of hydroxocobalamin and deferoxamine as antidotes. This discussion will explore the significance of our findings, address potential limitations, and propose directions for future research.

### Significance of Findings

Our research has demonstrated that cyanide, a potent cellular toxin, poses significant risks during fire incidents, particularly in environments where nitrogen-containing materials are present. The observed high lethality of cyanide at concentrations between 100–110 ppm in our smoke chamber experiments underscores the critical need for effective intervention strategies [14]. To establish a reliable model for generating high concentrations of hydrogen cyanide (HCN), we evaluated several combustion materials, including wood, polyurethane foam, acoustic cotton, and polypropylene (PP) plastic bags. Ultimately, melamine-based foam was selected due to its rapid release of high concentrations of HCN within a short combustion period, as confirmed by our preliminary tests using electrochemical sensors. Given the high risk associated with these experiments, all combustion tests were conducted within a fume hood equipped with appropriate protective gear. Additionally, we informed and obtained permission from the Environmental and Safety Center at National Yang Ming Chiao Tung University prior to initiating the experiments.

### Cyanide Quantification

We used hydrogen cyanide (HCN) aqueous solution testing reagents for cyanide quantification through colorimetry. This method is based on the color change produced by the reaction between cyanide and the reagent, which can be measured using a spectrophotometer. In colorimetric analysis, the cyanide reacts with the reagent to produce a colored reaction product. The absorbance of this product was measured at a specific wavelength (λmax = 580 nm).

To accurately determine the cyanide concentration in our simulated fire smoke environment, we implemented a method using colorimetric data to establish a calibration curve for cyanide analysis. This method provides a basis for precise measurement of cyanide concentrations, which is crucial for understanding its distribution and toxicity in fire smoke. In addition, this quantification approach was used to compare with the HCN detector readings as a secondary verification method, ensuring consistency and reliability in cyanide measurement.

### Calibration Curve Establishment

To determine cyanide concentrations, we first established a calibration curve. Known concentrations of cyanide standard solutions (ranging from 0 to 110 ppm) were reacted with the reagent, and their absorbance values were measured at λmax = 580 nm (Supple figure 5).

The absorbance values were plotted against the corresponding cyanide concentrations to generate the calibration curve. Linear regression analysis was used to calculate the slope and intercept of the calibration curve, establishing the relationship between concentration and absorbance.

### Sample Analysis

The collected cyanide aqueous solution was mixed with the reagent, and the absorbance value of the reaction product was measured.

Using the calibration curve, the measured absorbance value was converted to cyanide concentration. This method accurately determined the cyanide concentration in the simulated fire smoke environment.

### Data Processing

Each sample was measured at least three times to reduce error, and the average value was used.

The cyanide concentrations calculated from the colorimetric data were used to assess cyanide exposure levels in the fire smoke simulation environment. These data were further combined with survival experiment data from mice to analyze the relationship between cyanide concentration and its lethality.

By introducing colorimetric data for cyanide calibration, we achieved more precise measurements of cyanide concentrations. This precision is essential for understanding cyanide toxicity and evaluating the effectiveness of antidotes.

The significant improvement in survival rates and times in the drug-treated group compared to the control group provides strong evidence for the efficacy of hydroxocobalamin and deferoxamine in mitigating cyanide toxicity.

Hydroxocobalamin’s ability to bind cyanide and convert it into the non-toxic cyanocobalamin was evident in the reduced blood cyanide levels in treated mice (Supple figure 3, 4). Deferoxamine’s role in protecting against free radical damage further supports the combined use of these drugs as a comprehensive treatment approach.

### Recovery Rate Analysis

To better understand the operational losses and ensure accurate drug quantification, we incorporated recovery rate analyses as supplemental data [9, 10]. By using serum as the baseline and comparing recovery rates across formulations and nebulized samples, we identified potential degradation or deactivation of the drug. This approach allows us to validate the true concentrations of the nebulized drugs administered during the experiments, ensuring higher reliability in our efficacy assessments.

### Efficacy of Nebulized Drug Delivery

The use of nebulization to deliver hydroxocobalamin and deferoxamine directly to the respiratory tract proved highly effective. The significantly higher concentrations of these drugs in lung tissue compared to blood suggest efficient local delivery, which is crucial for rapid detoxification and protection of the respiratory system. This method offers practical advantages for emergency response scenarios, where quick and targeted intervention is essential.

### Potential Limitations

Despite the promising results, our study has several limitations. Firstly, the use of a controlled smoke chamber environment may not fully replicate the complexities and variabilities of real-world fire incidents. Factors such as varying smoke composition, environmental conditions, and the presence of other toxicants could influence the outcomes and should be considered in future studies.

Secondly, while mice serve as a relevant model for toxicological studies, there are physiological differences between mice and humans that may impact the generalizability of our findings. Further research involving other animal models or clinical trials would be necessary to validate the efficacy of these treatments in humans.

Thirdly, our study focused on the immediate effects of cyanide and the antidotes. Long-term health effects and potential delayed toxicities were not thoroughly examined. Extended observation periods and follow-up studies would be beneficial to understand any lasting impacts of cyanide exposure and treatment.

Future Research Directions

Building on our findings, several areas warrant further investigation:

Extended Toxicity Studies: Future research should explore the long-term health effects of cyanide exposure and the efficacy of antidotes over prolonged periods. This will provide a more comprehensive understanding of the chronic impacts and potential delayed toxicities.

Combination Therapies: While our study demonstrated the effectiveness of hydroxocobalamin and deferoxamine, investigating other potential antidotes or combination therapies could enhance treatment outcomes [15]. Exploring the synergistic effects of different drugs may offer improved protection against a broader range of toxicants in fire smoke [16,17, 18, 19].

Real-World Application: Conducting field studies or simulations that replicate real-world fire scenarios will help validate the practical applicability of our findings [20, 21]. Assessing the effectiveness of nebulized drug delivery in actual fire rescue operations can provide valuable insights for refining treatment protocols.

Advanced Detection Methods: Developing more sophisticated and rapid detection methods for cyanide and other toxicants in fire smoke could enhance early diagnosis and intervention. Portable and accurate detection devices would be beneficial for firefighters and emergency responders [17, 19].

Public Health and Safety Policies: Our research findings should inform the development of public health and safety policies aimed at reducing the risks associated with fire smoke inhalation [22]. Guidelines for the use of antidotes, training for emergency responders, and public awareness campaigns could significantly improve preparedness and response to fire incidents.

## Acknowledgments

The authors would like to thank the Instrumentation Center at National Yang Ming Chiao Tung University and Jin-Wu Tsai’s Lab for providing laboratory facilities and technical support, which were instrumental in completing this study. Histopathological analyses were performed by Yeong Jyi Chemical Apparatus Co., Ltd.

During the preparation of this work, the author(s) used ChatGPT to assist with referencing research and organizing the article. After utilizing this tool/service, the author(s) thoroughly reviewed and edited the content as needed and take full responsibility for the final publication.

## Concentrations of Hydroxocobalamin and Cyanocobalamin in Blood and Lung Samples Over Time

This table presents the measured concentrations of hydroxocobalamin and cyanocobalamin in blood and lung samples collected from C57BL/6 male mice after nebulized administration of two drug formulations [18]. The mice were divided into control and two experimental groups, receiving either 1X (25 mg/mL hydroxocobalamin + 1 mg/mL deferoxamine) or 5x (150 mg/mL hydroxocobalamin + 6 mg/mL deferoxamine), both reconstituted in 0.3M Tris buffer. Samples were collected at 20 minutes, 1 hour, and 2 hours post-administration. Data are expressed as mean ± standard deviation (ng/mL).

Mice received low dose, which included Hydroxocobalamin 25 mg and Deferoxamine 1 mg diluted in 0.3M Tris buffer, or high dose, which included Hydroxocobalamin 150 mg and Deferoxamine 6 mg diluted in 0.3M Tris buffer. Blood and lung fluid samples from smoke-exposed groups exhibited significantly higher Cyanocobalamin levels, supporting the conversion of Hydroxocobalamin to Cyanocobalamin in the presence of cyanide. Since each sample was processed independently, individual variability among mice may contribute to differences in drug concentrations. Additionally, the lung lavage process can result in variable drug recovery, which may lead to fluctuations in the measured concentrations. This table primarily serves to observe the impact of fire smoke on Hydroxocobalamin neutralization and its subsequent conversion to Cyanocobalamin.

Mean ± SEM values are presented for each condition (n = 3 per group). Statistical analysis was performed using a one-way ANOVA to compare Hydroxocobalamin and Cyanocobalamin concentrations across different groups, followed by a post-hoc Tukey’s test to identify significant pairwise differences. Significance was set at p < 0.05, with higher Cyanocobalamin levels observed in smoke-exposed groups compared to lung fluid samples.

## Supplemental data

S1 Figure. **Nebulization and Animal Grouping**

The study utilized 8–10-week-old B6(C57BL/6) mice, assigned to receive either a low dose (25 mg/mL hydroxocobalamin + 1 mg/mL deferoxamine) or a high dose (150 mg/mL hydroxocobalamin + 6 mg/mL deferoxamine) via nebulization. Both formulations were reconstituted in 0.3 M Tris buffer. A minimum of three mice per group was included to ensure statistical reliability. Following inhalation, the mice were observed over an one-week period to assess therapeutic efficacy and toxicological safety. Throughout the observation period, none of the mice displayed adverse reactions, abnormal behavior, or mortality, demonstrating the tolerability of both dosing regimens.

S2_Figure_**Mortality Distribution in Mice Exposed to HCN**

A mortality distribution analysis was conducted to examine the relationship between hydrogen cyanide (HCN) concentration, exposure duration, and survival rate in mice. Figure 1 presents the mortality distribution across different HCN concentrations and exposure times, with each data point representing a unique concentration-duration combination. The distribution highlights the dose-dependent toxicity of HCN, where increased concentrations and prolonged exposure durations correlate with higher mortality rates. This distribution provides critical insights into the lethal thresholds of HCN, helping to establish safe exposure limits and improve understanding of acute toxicity mechanisms.

S3 Figure. **Calibration curve of Hydroxocobalamin**

The compound analyzed was hydroxocobalamin, with a correlation coefficient of r = 0.998196 and r^2^ = 0.996395. The calibration curve equation was 14.3037 * x + - 15.7776, using an external standard response type based on area. The curve type was linear, with the origin included, and the weighting was 1/x, with no axis transformation applied.

S4 Figure. **Calibration curve of Cyanocobalamin**

The compound analyzed was cyanocobalamin, with a correlation coefficient of r = 0.996805 and r^2^ = 0.993620. The calibration curve equation was 30.8594 * x + - 9.71672, using an external standard response type based on area. The curve type was linear, with the origin included, and the weighting was 1/x, with no axis transformation applied.

S5 Figure. **Calibration curve for hydrogen cyanide (HCN) concentration determination**

The curve was obtained through quadratic regression analysis of measured HCN concentrations (ppm vs. mg/L) using colorimetric assays. The fitted equation is y = 0.00022x^2^ - 0.0536x + 3.2820, with a high correlation coefficient (R^2^) of 0.9993, indicating excellent fit quality and reliability for quantitative analyses.

S6 Figure. **Lung histopathological changes after smoke inhalation and preventive drug administration**

Representative hematoxylin and eosin (H&E) stained lung tissue sections under 4x (upper panel, scale bar: 1000 μm) and 20x magnification (lower panel, scale bar: 200 μm). Groups shown include Control (no smoke exposure), Smoke (smoke exposure only), Drug+Smoke (Low dose preventive treatment followed by smoke exposure), and Drug+Smoke (High dose preventive treatment followed by smoke exposure).

S7 Figure. **Hepatic histopathological evaluation after drug inhalation at high dosage** Representative liver sections stained with hematoxylin and eosin (H&E) under 4x (upper panel, scale bar: 1000 μm) and 20x magnification (lower panel, scale bar: 200 μm). The figure compares Control (non-treatment) and Drug High-dose groups (tissue collected 20 minutes after drug inhalation), showing no observable pathological changes in hepatic tissue upon drug administration.

S1 Table. **Baseline Serum Recovery Rate**

This table shows the recovery rates of the drug using serum as the baseline. The data highlights potential losses during the operational process, including degradation or deactivation of the drug.

S2 Table **Recovery Rate for 25 mg/mL Hydroxocobalamin + 1 mg/mL Deferoxamine**

This table demonstrates the recovery rates observed when using the low-dose formulation. It provides insights into the stability and potential degradation patterns of the nebulized drug under the experimental conditions.Blood: Refers to samples collected from blood serum. Lung: Refers to samples collected from lung tissue or lung fluid.

S3 Table. **Recovery Rate for 150 mg/mL Hydroxocobalamin**

This table displays the recovery rates for the high-dose formulation. By comparing this with the baseline serum recovery rate, one can infer the true concentrations achieved post-nebulization and the operational losses during the process. Blood: Refers to samples collected from blood serum. Lung: Refers to samples collected from lung tissue or lung fluid. SL: Refers to lung fluid samples collected from lung tissues after smoke exposure.

## Concentrations of Hydroxocobalamin and Cyanocobalamin in Blood and Lung Samples Over Time

This table presents the measured concentrations of hydroxocobalamin and cyanocobalamin in blood and lung samples collected from C57BL/6 male mice after nebulized administration of two drug formulations (18). The mice were divided into control and two experimental groups, receiving either 1X (25 mg/ml hydroxocobalamin + 1 mg/ml deferoxamine) or 5x (150 mg/ml hydroxocobalamin + 6 mg/ml deferoxamine), both reconstituted in 0.3M Tris buffer. Samples were collected at 20 minutes, 1 hour, and 2 hours post administration. Data are expressed as mean ± standard deviation (ng/ml).

Mean+ SEM values are presented for each condition (n = 3 per group).Statistical analysis was performed using a one-way ANOVA to compare Hydroxocobalamin and Cyanocobalamin concentrations across different groups, followed by a post-hoc Tukey’s test to identify significant pairwise differences. Significance was set at p < 0.05, with higher Cyanocobalamin levels observed in smoke-exposed groups compared to lung fluid samples.

